# Cbln1 directs axon targeting by corticospinal neurons specifically toward thoraco-lumbar spinal cord

**DOI:** 10.1101/2022.04.06.487184

**Authors:** Janet H.T. Song, Carolin Ruven, Payal Patel, Frances Ding, Jeffrey D. Macklis, Vibhu Sahni

## Abstract

Corticospinal neurons (CSN) are centrally required for skilled voluntary movement, which necessitates that they establish precise subcerebral connectivity with the brainstem and spinal cord. However, molecular controls regulating specificity of this projection targeting remain largely unknown. We previously identified that developing CSN subpopulations exhibit striking axon targeting specificity in the spinal white matter. These CSN subpopulations with segmentally distinct spinal projections are also molecularly distinct; a subset of differentially expressed genes between these distinct CSN subpopulations function as molecular controls regulating differential axon projection targeting. Rostrolateral CSN extend axons exclusively to bulbar-cervical segments (CSN_BC-lat_), while caudomedial CSN (CSN_medial_) are more heterogeneous, with distinct, intermingled subpopulations extending axons to either bulbar-cervical or thoraco-lumbar segments. Here, we report that *Cerebellin 1* (*Cbln1*) is expressed specifically by CSN in medial, but not lateral, sensorimotor cortex. *Cbln1* shows highly dynamic temporal expression, with *Cbln1* levels in CSN highest during the period of peak axon extension toward thoraco-lumbar segments. Using gain-of-function experiments, we identify that Cbln1 is sufficient to direct thoraco-lumbar axon extension by CSN. Mis-expression of Cbln1 in CSN_BC-lat_ either by *in utero* electroporation, or in postmitotic CSN_BC-lat_ by AAV-mediated gene delivery, re-directs these axons past their normal bulbar-cervical targets toward thoracic segments. Further, Cbln1 overexpression in postmitotic CSN_medial_ increases the number of CSN_medial_ axons that extend past cervical segments into the thoracic cord. Collectively, these results identify that Cbln1 functions as a potent molecular control over thoraco-lumbar CSN axon extension, part of an integrated network of controls over segmentally-specific CSN axon projection targeting.

**Significance Statement:** Corticospinal neurons (CSN) exhibit remarkable diversity and precision of axonal projections to targets in the brainstem and distinct spinal segments; the molecular basis for this targeting diversity is largely unknown. CSN subpopulations projecting to distinct targets are also molecularly distinguishable. Distinct subpopulations degenerate in specific motor neuron diseases, further suggesting that intrinsic molecular differences might underlie differential vulnerability to disease. Here, we identify a novel molecular control, Cbln1, expressed by CSN extending axons to thoraco-lumbar spinal segments. Cbln1 is sufficient, but not required, for CSN axon extension toward distal spinal segments, and *Cbln1* expression is controlled by recently identified, CSN-intrinsic regulators of axon extension. Our results identify that Cbln1, together with other regulators, coordinates segmentally precise CSN axon targeting.

## Introduction

For skilled motor control, the cerebral cortex must precisely and accurately connect with specific spinal segments (Sahni et al., 2020). How corticospinal neuron (CSN) axonal projection targeting is established during development underlies motor function, CNS organization, and species differences in orofacial and forelimb dexterity during evolution. Prior work in the field has identified molecular controls over CSN specification, development, and projection targeting (Arlotta et al., 2005; Molyneaux et al., 2005; Chen et al., 2005; Pang et al., 2000b; Lai et al., 2008; Chen et al., 2008; Kwan et al., 2008; Joshi et al., 2008; Tomassy et al., 2010; McKenna et al., 2011; Han et al., 2011; Shim et al., 2012; Lodato et al., 2014; Muralidharan et al., 2017; Diaz et al., 2020; Sahni et al., 2021a,b).

We recently identified that developing CSN subpopulations exhibit striking axon targeting specificity in the spinal white matter, and that this establishes the foundation for durable specificity of adult corticospinal circuitry. CSN_BC-lat_, which reside in rostro-lateral cortex, are relatively homogeneous, with projections to only bulbar-cervical segments. In contrast, CSN residing in medial sensorimotor cortex (CSN_medial_) are more heterogeneous, with distinct interdigitated subpopulations extending axons to either bulbar-cervical or thoraco-lumbar segments– CSN_BC-med_ extend axons only to bulbar-cervical segments, while CSN_TL_ extend axons past cervical cord to thoracic and lumbar spinal segments. We further identified that these segmentally distinct CSN subpopulations are molecularly distinct from early development, enabling molecular delineation and prospective identification even before eventual axon-targeting decisions are evident in the spinal cord: 1) *Klhl14* expression delineates *Klhl14*-positive CSN_BC-lat_ from *Klhl14*-negative CSN_BC-med_; 2) all CSN_TL_ are *Klhl14*-negative; and 3) nearly all CSN_TL_ express *Crim1* (schematized in Fig. 1A) (Sahni et al., 2021a).

**Figure 1:**
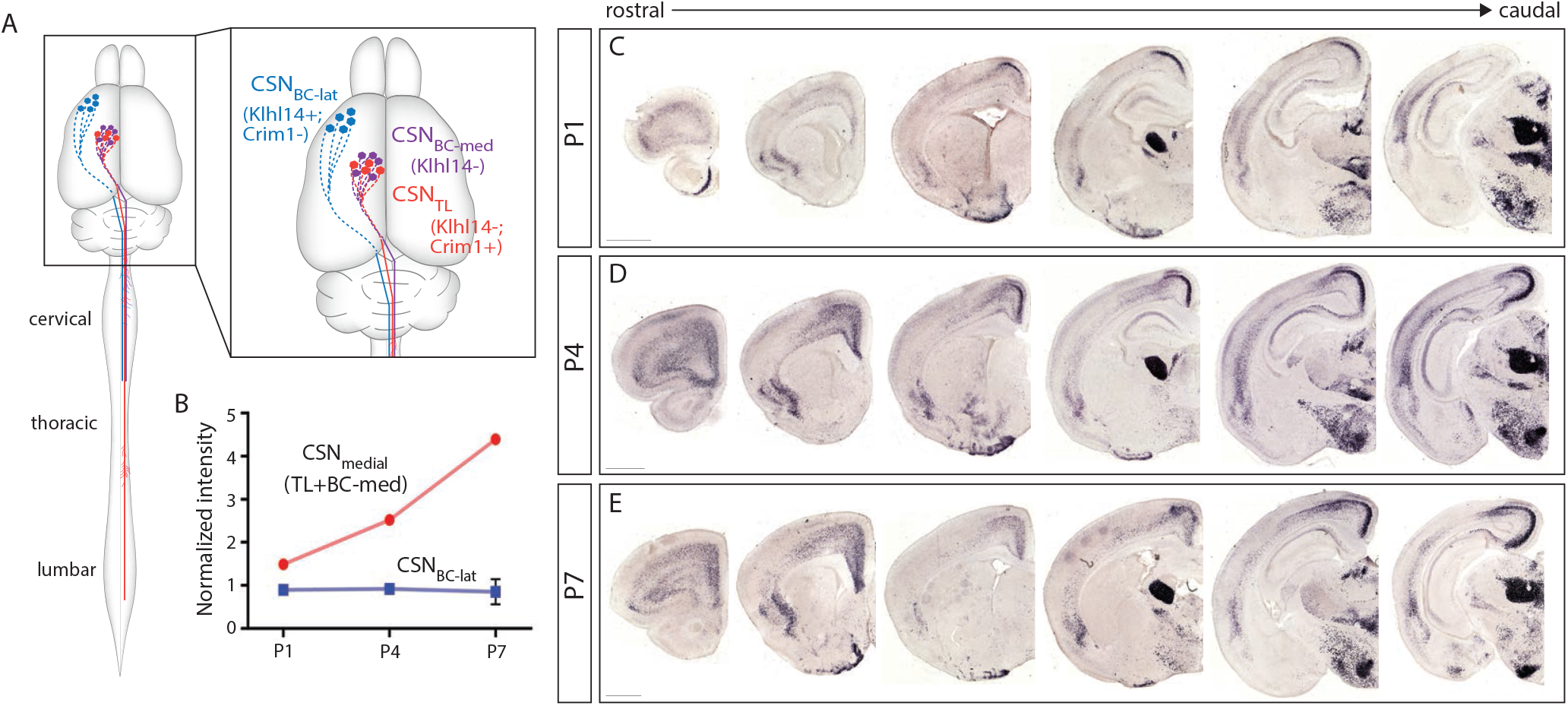
Cbln1 is specifically expressed by CSN residing in medial sensorimotor cortex. **(A)** Schematic of the mouse brain and spinal cord, with inset delineating the three spatially, segmentally, and molecularly distinct CSN subpopulations: CSN_BC-lat_ (blue) reside in rostro-lateral sensorimotor cortex and extend axons only to bulbar-cervical segments; CSN_TL_ (red) reside in medial sensorimotor cortex and extend axons to thoraco-lumbar spinal segments; and CSN_BC-med_ (purple) also reside in medial sensorimotor cortex and extend axons only to bulbar-cervical segments. CSN_TL_ and CSN_BC-med_ are both located in medial sensorimotor cortex, cannot be spatially distinguished, and are collectively referred to as CSN_medial_. *Klhl14* expression delineates *Klhl14*-positive CSN_BC-lat_ from *Klhl14*-negative CSN_medial_. Nearly all CSN_TL_ express *Crim1* while CSN_BC-lat_ are *Crim1*-negative. **(B)** Prior gene expression analysis of CSN_BC-lat_ and CSN_medial_ identified *Cbln1* as a gene that is not expressed by CSN_BC-lat_ (blue) but whose expression increases from P1 to P7 in CSN_medial_ (red) (Sahni et al., 2021a). **(C)** *In situ* hybridization confirms that *Cbln1* is expressed in layer V, where CSN reside. *Cbln1* expression increases from P1 to P7 and is restricted to medial layer V throughout the rostrocaudal extent of sensorimotor cortex. Scale bars are 1mm.

Crim1 and Klhl14 direct differential CSN axon segmental targeting by these subpopulations (Sahni et al., 2021b), indicating that the diversity of CSN axonal targeting is controlled in part by CSN-intrinsic mechanisms. Crim1 is both necessary and sufficient for CSN_TL_ axon extension to thoracic and lumbar segments. Crim1 mis-expression is sufficient to re-direct a subset of CSN_BC-lat_ axons into the caudal thoracic cord. However, this effect of Crim1 mis-expression, though striking, only affects a minority of the overall CSN_BC-lat_ subpopulation, with the majority of CSN_BC-lat_ axons terminating in the cervical cord. In addition, although a subset of CSN_TL_ axons, which normally extend past the cervical cord, fail to extend to caudal thoracolumbar segments in *Crim1* null mice, ∼ 50% of CSN_TL_ axons still reach the lumbar cord. This indicates that some CSN_TL_ axons can extend to distal spinal targets independent of Crim1 function. Collectively, these results indicate that there are likely additional regulators that direct CSN_TL_ axon extension to distal spinal segments.

Here, we identify Cbln1 as a novel regulator of CSN axon targeting to thoraco-lumbar spinal segments. Cbln1 is a member of the C1q superfamily, which includes proteins critically essential for normal function of the immune and nervous systems (Ghai et al., 2007; Stevens et al., 2007; Yuzaki, 2011). Cbln1 has been extensively characterized in the cerebellum, where it is required for synapse formation and synapse stabilization between parallel fibers of granule cells and Purkinje cell dendrites (Hirai et al., 2005; Matsuda et al., 2010; Uemura et al., 2010; Elegheert et al., 2016; Ibata et al., 2019; Takeo et al., 2021). More recently, it also has been shown to play roles in synapse formation in the striatum and the hippocampus (Kusnoor et al., 2010; Seigneur and Südhof, 2018). There is no previously reported function for Cbln1 in corticospinal connectivity.

We find that *Cbln1* is expressed specifically by CSN_medial_, and its expression coincides with the peak period of CSN axon extension toward thoraco-lumbar spinal segments. Misexpression of Cbln1 in CSN_BC-lat_ is sufficient to re-direct axons past their normal targets in the cervical cord toward distal thoracic segments. We also identify that Cbln1 over-expression in CSN_medial_ can increase the number of axons extending past the cervical cord toward thoraco-lumbar segments. This suggests that Cbln1 is sufficient to direct thoraco-lumbar extension by CSN axons in these contexts. Further, this effect on CSN axon extension occurs prior to axon collateralization and synapse formation, establishing a novel function for Cbln1 in directing axon extension, independent of its known functions in synapse formation established elsewhere in the central nervous system. Together, these results identify Cbln1 as a novel, CSN-intrinsic determinant of thoraco-lumbar segmental axon targeting specificity.

## Materials and Methods

### Mice

CD-1 mice (Charles River Laboratories, Wilmington, MA) were used for gene expression analysis, *in utero* electroporation, and AAV injections. *Fezf2* null mice were generated previously (Hirata et al., 2004) and have been described (Molyneaux et al., 2005). *Cbln1* null mice were generated and described previously (Hirai et al., 2005). E0.5 was set as the day of the vaginal plug, and P0 was set as the day of birth. Mice received food and water *ad libitum*, and were housed on a 12-hour on/off light cycle. All mouse studies were approved by the IACUC at Harvard University and at Weill Cornell Medicine. All studies were performed in accordance with institutional and federal guidelines.

### Tissue collection and preparation

Mice were anesthetized by hypothermia (P0-P4) or with an intraperitoneal injection (P7-adult) of 0.015 mL/g body weight Avertin (1.25% 2-2-2 tribromoethanol in a solvent that contains 0.63% isoamyl alcohol by weight in ddH_2_O). Mice were perfused transcardially, first with PBS then with 4% paraformaldehyde (PFA) for fixation. The skull, musculature, limbs, and internal organs (viscera) were removed from the thoracic and abdominal cavities. The remaining skeletal structures were postfixed overnight in 4% PFA at 4°C. The brains were also dissected out, and postfixed overnight in 4% PFA at 4°C. The following day, the spinal cords were dissected out of the vertebral column. The brains and spinal cords were then washed in 1X PBS, and stored in 1X PBS at 4°C.

To collect embryonic tissue (E18.5), timed pregnant females were anesthetized with an intraperitoneal injection of 1 mL Avertin, and euthanized with an additional 1 mL intracardiac injection of Avertin. Embryos were dissected from the uterine horn and decapitated, and the entire head was fixed overnight in 4% PFA at 4°C. The following day, the brains were dissected out, washed in 1X PBS, and stored in 1X PBS at 4°C.

For immunocytochemistry and *in situ* hybridization, brains or spinal cords were placed in Tissue-Tek OCT Compound (Sakura Finetek, Torrance, CA) for sectioning using a cryostat (Leica CM3050 S, Wetzlar, Germany). Prior to sectioning, the cerebellum, pons, and medulla were removed with a razor blade, leaving the forebrain. 50 μm coronal brain sections, or 50 μm axial or sagittal spinal cord sections, were obtained using a cryostat. All sections were stored in 1X PBS at 4°C.

### In situ hybridization and immunocytochemistry

*In situ* hybridization was performed as previously described (Arlotta et al., 2005). The primer sequences used to generate the *in situ* hybridization probes are from the Allen Brain Atlas (http://www.brain-map.org). Brains were fixed and stained using standard methods (Molyneaux et al., 2005), with the primary antibody rabbit anti-GFP, 1:500 (Invitrogen).

### Retrograde labeling of corticospinal neurons

CSN that project to lumbar spinal segments were retrogradely labeled at P5 with an Alexa Fluor 555-conjugated cholera toxin subunit B (CTB-555) recombinant retrograde tracer (Invitrogen). For these injections, mice were anesthetized under ice for 4 minutes, then visualized by Vevo 770 ultrasound backscatter microscopy (VisualSonics, Toronto, Canada) using Aquasonic 100 ultrasound gel (Parker Laboratories, Fairfield, NJ). 4 slow-pulse injections of 60 nl of CTB-555 (2 mg/ml) were deposited on each side of the midline at L1-L2 using a pulled glass micropipette with a nanojector (Nanoject II, Drummond Scientific, Broomall, PA) to obtain bilateral labeling. The mice were placed on a heating pad for recovery. Mice were euthanized at P7, allowing the retrograde tracers 2 days for transport.

### Anterograde labeling of corticospinal neurons

P28 mice were anesthetized using isofluorane anesthesia (2.5% in 100% O_2_), and injected in caudomedial cortex with the anterograde tracer biotinylated dextran amine (BDA) using the following stereotactic coordinates - 1.0 mm lateral to the midline at Bregma at a depth of 0.8 mm. A glass capillary micropipette was filled with a 10% solution of BDA (10,000 MW; Thermo Fisher) which was delivered into the cortex by iontophoresis using constant current conditions (8 microamps; 7 seconds on 7 seconds off) for a total period of 20 minutes. Mice were perfused at P35. The injection site and labeled axons were visualized using DAB staining (Vector Laboratories, Burlingame, CA).

### In utero electroporation

Surgeries were performed as previously described (Molyneaux et al., 2005; Greig et al., 2016). To generate the *Cbln1* over-expression construct, *Cbln1* cDNA was cloned 3’ to an EGFP coding sequence, which was driven by the CAG promoter, and the two ORFs were separated by the t2A linker sequence. In the control plasmid, the *Cbln1* cDNA was replaced with a STOP codon 3’ to the t2A linker sequence.

### AAV-mediated gene delivery

Constructs expressing GFP or *Cbln1* were packaged into AAV 2/1, a serotype known to be specific to neuronal expression, by the Massachusetts General Hospital Virus Core using established protocols. At P0, the appropriate viral mixture (103.5 ng of AAV, 0.05% DiI, and 0.08% Fast Green in 1X PBS) was injected at 23 nl per injection into specific cortical subregions using the same set-up described previously for *in utero* electroporation (Molyneaux et al., 2005; Greig et al., 2016). All viral work was approved by the Harvard Committee on Microbiological Safety, and the Institutional Biosafety Committee at Weill Cornell Medicine; all work was conducted according to institutional guidelines.

### Imaging and quantification

For all CST quantification on axial sections, 60X confocal Z stacks of the entire CST in the dorsal funiculus were obtained on either a Biorad Radiance 2100 confocal microscope (Biorad, Hercules, CA), a Zeiss LSM 880 confocal microscope (Zeiss, Oberkochen, Germany), or a Leica SP8 confocal microscope (Leica Microsystems). Cervical, thoracic, and/or lumbar cord axial sections were imaged using identical parameters. 4X and 10X images of brain and spinal cord sections were obtained on either an ANDOR Clara DR328G camera (ANDOR Technology, South Windsor, CT) mounted on a Nikon Eclipse 90i microscope (Nikon Instruments) or a Zeiss Axioimager M2 (Zeiss) using Stereo Investigator software (MBF Biosciences). For counts of CSN retrogradely labeled from lumbar L1L2, we first examined the spinal cord to confirm the matched spinal level of the retrograde tracer injection. Following this confirmation, we imaged and analyzed matched coronal sections for each of three specific rostrocaudal levels in wild-type, *Cbln1* heterozygous, and *Cbln1* null mice. *Cbln1* wild-type and *Cbln1* heterozygous mice are indistinguishable in this line (Hirai et al., 2005). The labeled neurons were counted using the cell counter function in ImageJ (National Institutes of Health, Bethesda, MD). In all mice, regardless of the genotype, labeled neurons were only found in the medial cortex upon retrograde injection in L1-L2.

For counts of anterogradely labeled, BDA+ axons in the cervical, thoracic, and lumbar spinal cords, we first examined coronal sections of the brain to confirm matched sites of anterograde tracer injection. We then counted the number of axons present at the cervical C1-C2, thoracic T1-T2, and lumbar L1-L2 levels using the cell counter function in ImageJ.

For EGFP+ axon counts in axial sections, 3 axial sections were imaged at both cervical C1-C2 and thoracic T1-T2 levels per mouse, and the axon counts were averaged from 3 separate sections. For axon intensity measurements in axial sections, at least 3 axial sections were imaged for each mouse at cervical C1-C2, thoracic T1-T2, and lumbar L1-L2. Background fluorescence intensity was measured from the maximum intensity projection of each Z stack image, and subtracted from the Z stack using ImageJ, such that the intensity of parts of the section without labeled axons was zero. The dorsal funiculus was then selected as the region of interest, and the intensity was measured in each Z stack image. The top 3 measurements from the Z stack were averaged as the fluorescence intensity for that section. These measurements were then averaged for at least 3 axial sections at each spinal cord segmental level.

For axon extension experiments, the thoracic cord was sectioned sagittally, and every section that contained a labeled axon was imaged. Each such section was imaged in its entirety, from rostral to caudal and throughout the medio-lateral Z axis. Z stacks for each such section were collapsed to a single two-dimensional plane using the “create focused image” function on the NIS-Elements acquisition software (Nikon Instruments). The collapsed sections were then further combined into one two-dimensional image per mouse by aligning the edges of each section and performing a maximum intensity projection across all sections in Adobe Photoshop using the “Lighten” mode with 100% opacity. This single two-dimensional image per mouse was converted into a monochrome image. To quantify axon extension, we cropped the single image per mouse into dorso-ventral rectangular regions at five rostro-caudal locations (rostral-most, 25% caudal, 50% caudal, 75% caudal, and caudal-most). In each such rectangular region, the CST was then selected as the region-of-interest, and fluorescence intensity was measured in ImageJ. Background fluorescence intensity was measured at an immediately adjacent location in the image and subtracted from this measurement. Intensity at each rostrocaudal level was then normalized to the intensity at the rostral-most limit of the thoracic cord. If no labeled axons were present at the rostrocaudal limit in an individual case, the intensity was set to 0 in that case.

For all of the experiments, the experimenter analyzing the images remained blinded to the experimental conditions.

### Experimental design and statistical analysis

Data are presented as mean ± SEM (standard error of mean), with *n* indicating the number of mice used in each group for comparison. The Student’s T-test was used to assess whether the proportion of CSN axons that reach thoracic T1-T2 from cervical C1-C2 or that reach lumbar L1-L2 from cervical C1C2 was significantly different between mice injected with GFP (control) or Cbln1. We used the two-tailed Student’s T test for the *in utero* electroporation experiment, then used the one-tailed Student’s T test in the AAV injection experiments to test the hypothesis that *Cbln1* promotes axon extension into the thoracic spinal cord. To model the distribution of the proportion of axons that reach T1-T2 from cervical C1-C2 (T1/C1) following injection with either control AAV-EGFP or AAV-Cbln1, we fit a mixture model of two Gaussians using the R package *mixtools*. The two-tailed Student’s T test was also used to compare the proportion of retrogradely-labeled neurons at distinct rostral-caudal levels when comparing WT and *Cbln1* null mice. We used a two-way ANOVA with repeated measures followed by Fisher’s least significant difference posthoc test for the axon extension analyses. Data distribution was assumed to be normal, but this was not formally tested. Male and female mice were used without distinction in experiments.

## Results

### Cbln1 is expressed by CSN in medial, but not lateral, sensorimotor cortex during early postnatal development

We previously performed differential gene expression analysis to identify potential candidate molecular controls over CSN axon targeting to bulbar-cervical versus thoraco-lumbar spinal segments (Sahni et al., 2021a). We compared gene expression at three critical timepoints, P1, P4, and P7, between CSN in lateral sensorimotor cortex (CSN_BC-lat_), which extend projections exclusively to bulbar-cervical spinal segments, and CSN in medial sensorimotor cortex (CSN_medial_), with two interspersed and molecularly distinct subpopulations that extend projections to both bulbar-cervical (CSN_BC-med_) and thoraco-lumbar (CSN_TL_) segments (schematized in Fig. 1A). By P4, CSN_TL_, in contrast to CSN_BC-lat_ and CSN_BC-med_, extend axon projections toward distal spinal cord segments (Bareyre et al., 2005; Kamiyama et al., 2015; Sahni et al., 2020, 2021a). To identify candidate molecular controls that control CSN_TL_ axonal targeting specifically within CSN_medial_, we previously investigated genes that exhibit significant differential expression between CSN_BC-lat_ and CSN_medial_ at P4 (Sahni et al., 2021a,b). This work identified multiple candidate molecular regulators of axon targeting and collateralization, including *Crim1* as both a unique identifier of CSN_TL_, and a molecular control that is both necessary and sufficient to direct CSN axon extension to thoraco-lumbar segments. This further validated the original differential gene expression analyses to identify critical regulators over segmentally-specific CSN axon targeting.

Another candidate molecular control identified by this approach is *Cbln1*, which is specifically expressed by CSN_medial_ but not by CSN_BC-lat_, with peak differential expression at P4 and P7 (Fig. 1B). This peak coincides with the period when CSN_TL_ axons extend past the cervical cord toward thoraco-lumbar segments. This time course of significant differential expression along with known roles for Cbln1 and its family members in synaptogenesis (Hirai et al., 2005; Stevens et al., 2007; Kusnoor et al., 2010; Matsuda et al., 2010; Uemura et al., 2010; Seigneur and Südhof, 2018; Ibata et al., 2019; Takeo et al., 2021) suggested that Cbln1 might function in controlling these processes by some or all CSN_medial_.

To investigate this hypothesis, we first confirmed these transcriptomic data by investigating *Cbln1* expression in the developing sensorimotor cortex using *in situ* hybridization. We examined expression at the three developmental times analyzed by differential gene expression analysis– P1, P4, and P7. These experiments identify that, at all three developmental times, *Cbln1* is expressed in medial, but not lateral, layer V (Fig. 1C-E). *Cbln1* expression remains restricted to medial layer V throughout the rostro-caudal extent of the sensorimotor cortex (Fig. 1C-E).

To determine whether *Cbln1* is specifically expressed by CSN in layer V, we investigated *Cbln1* expression in *Fezf2* null mice, which completely lack CSN (Chen et al., 2005; Molyneaux et al., 2005). At P7, *Cbln1* expression in layer V is completely abolished in *Fezf2* null mice, confirming that *Cbln1* is expressed specifically by CSN in layer V (Fig. 2). The abolition of *Cbln1* expression in medial layer V in *Fezf2* null mice was also observed at P1 and P4 (data not shown). As expected, *Cbln1* expression in layers II/III and VI remains unchanged in *Fezf2* null mice (Fig. 2). This specific expression of *Cbln1* by CSN_medial_ throughout early postnatal development suggests that Cbln1 might function in a CSN subpopulation-specific manner.

**Figure 2:**
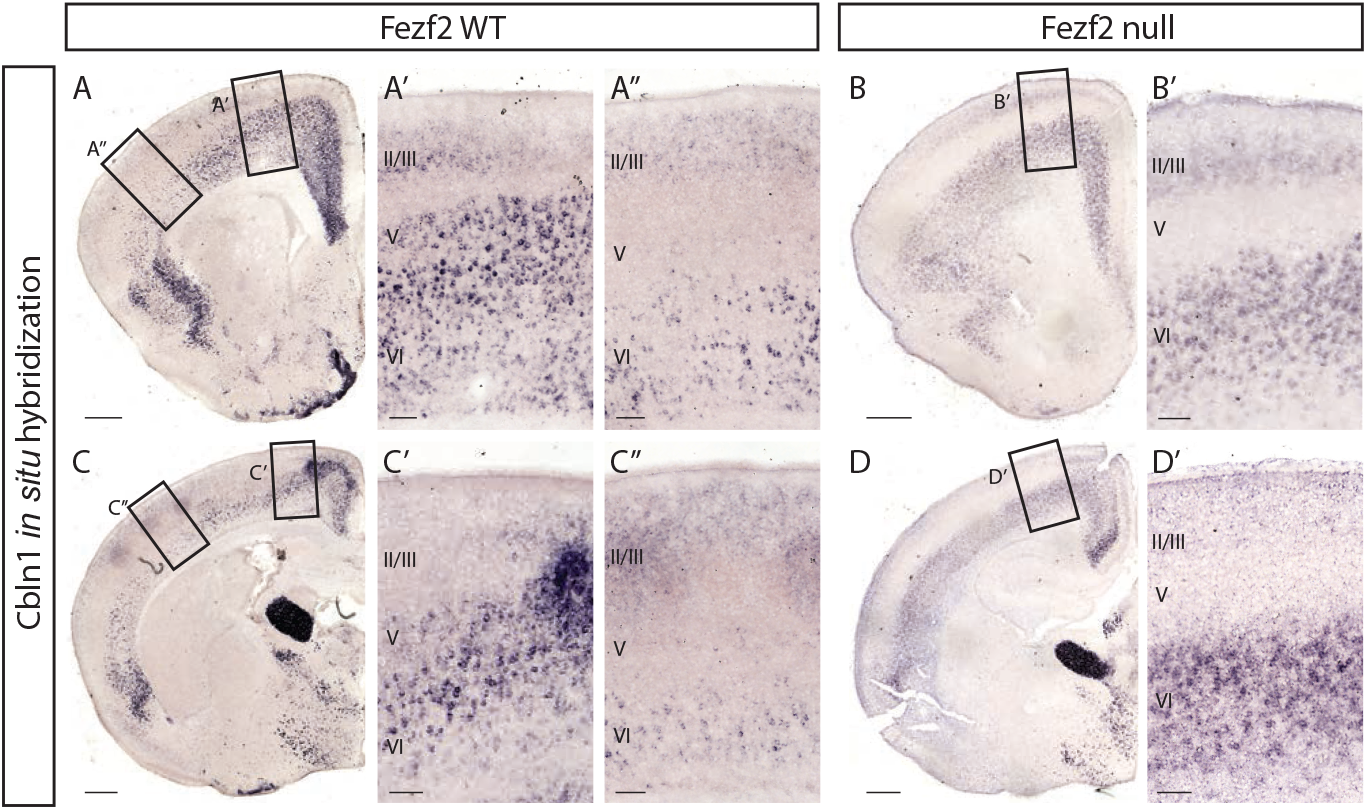
Cbln1 is specifically expressed by CSN in layer V. *In situ* hybridization at P7 shows that *Cbln1* is expressed in medial layer V in *Fezf2* WT (**A**,**C**) but not in *Fezf2* null (**B**,**D**) mice. *Fezf2* null mice completely lack CSN (Molyneaux et al., 2005; Chen et al., 2005). This indicates that *Cbln1* is expressed only by CSN in layer V in medial sensorimotor cortex. Scale bars are 100 μm for insets and 500 μm for other images.

### Cbln1 is expressed in neocortical and subcortical regions during early postnatal development

To rigorously investigate the spatial and temporal course of *Cbln1* expression in sensorimotor cortex from development into maturity, we performed *in situ* hybridization for *Cbln1* in WT mice at E18.5, P1, P4, P7, P10, P14, P21, P28, and adult (>3 month old). At E18.5, *Cbln1* is not expressed in the sensorimotor cortex (data not shown), indicating that Cbln1 is not required for early CSN development. At P1, *Cbln1* is expressed in medial layer V throughout the rostral-caudal extent of sensorimotor cortex (Fig. 3A,B). *Cbln1* expression then steadily increases in medial layer V from P4 to P7 (Fig. 2A,C, Fig. 3C,D). This expression decreases slightly at P10 (Fig. 3E,F). *Cbln1* is expressed at very low levels in layer V by P14, and is absent in layer V by P21 (Fig. 3G-K). *Cbln1* expression was never observed in lateral layer V at any time point (Fig. 3). These results confirm the initial observations of differential *Cbln1* expression by CSN_medial_ versus CSN_BC-lat_ (Sahni et al., 2021a). Further, these results also highlight the temporal dynamics of *Cbln1* expression with peak expression around P7 and declining expression levels thereafter.

**Figure 3:**
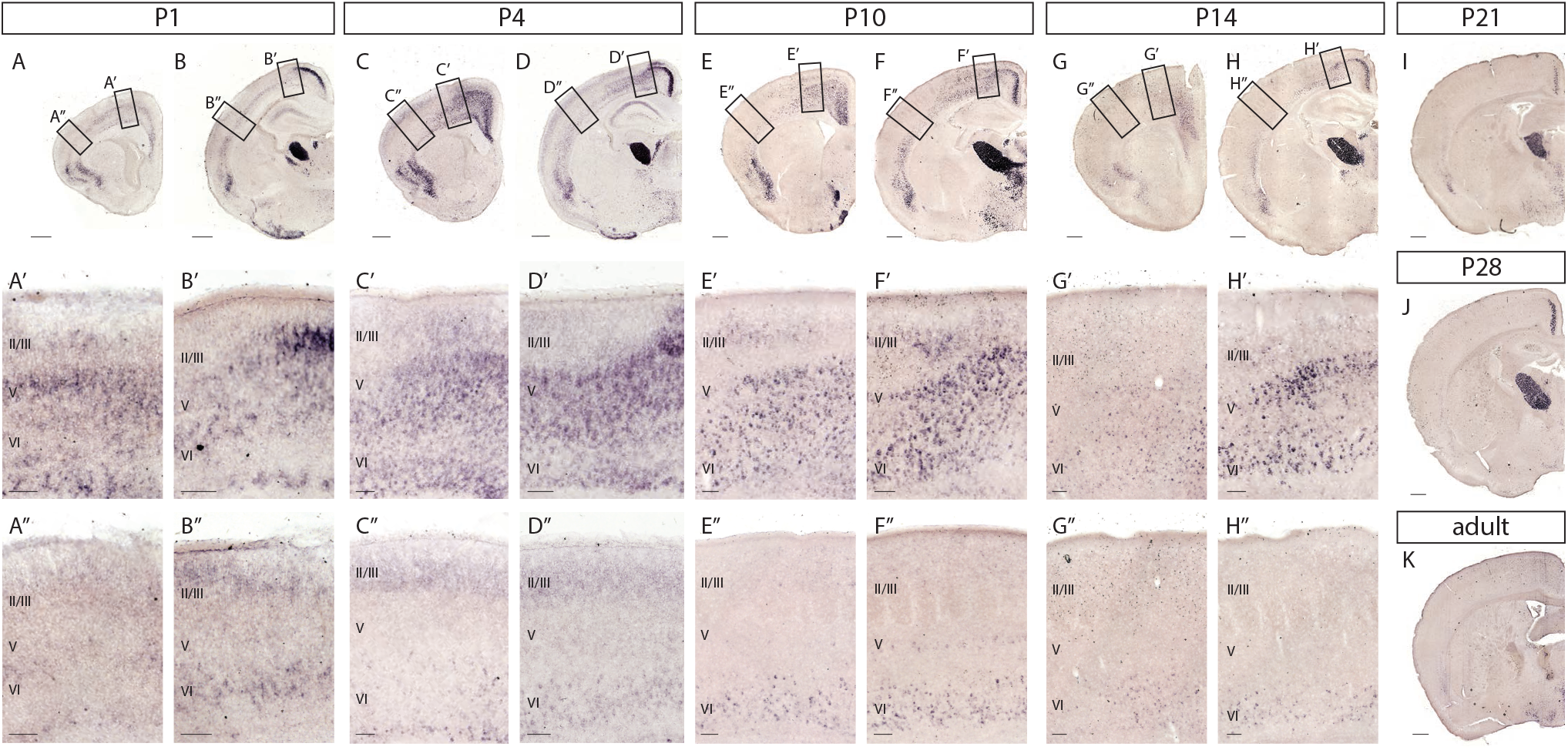
Time course of *Cbln1* expression. *Cbln1* is expressed throughout the rostro-caudal extent of sensorimotor cortex in medial but not lateral layer V at P1 **(A**,**B)**, P4 (**C**,**D**), P10 (**E**,**F**), and P14 (**G**,**H**). *Cbln1* expression in layer V is absent in P21 **(I)**, P28 **(J)**, and in >3 month old **(K)** mice. Scale bars are 100μm for insets and 500μm for all other images.

Within the cortex, *Cbln1* is also expressed in other layers in the neocortex and in subcerebral regions during postnatal development. In layer VI, *Cbln1* is expressed at low levels rostrolaterally. *Cbln1* expression in layer VI increases in caudomedial sensorimotor cortex from P1 until P10, and is present at low levels by P14 (Fig. 2, Fig. 3). *Cbln1* is expressed in medial layers II/III, with a higher expression level in caudal versus rostral sensorimotor cortex (Fig. 3). *Cbln1* is also highly expressed in the cingulate and piriform cortex from P1 to P28, with expression decreasing with age (Fig. 3). Consistent with previous reports (Iijima et al., 2007; Kusnoor et al., 2010; Otsuka et al., 2016), we find *Cbln1* is also very highly expressed outside the neocortex - in the thalamus and the hypothalamus from E18.5 into adulthood, and in the septum at P10 and P14 (Fig. 3).

Although *Cbln1* is expressed in many neocortical and subcerebral regions, its specific restriction within layer V to medial sensorimotor cortex combined with its distinct time course of expression suggests that Cbln1 might perform CSN_TL_-specific functions. Interestingly, *Cbln1* expression in layer V is largely confined to early postnatal development, with expression increasing from P1 to P7 and then decreasing by P14 (Fig. 2, Fig. 3). This corresponds with the time course of CSN_TL_ axon extension to caudal spinal segments during development. CSN_TL_ first extend axons toward the lumbar cord by P5, with the number of axons reaching the lumbar cord steadily increasing from P7 to P14, with continued extension up to P28 (Bareyre et al., 2005; Kamiyama et al., 2015; Sahni et al., 2020, 2021a). The restriction of *Cbln1* expression to medial CSN with peak expression coincident with the time period of CSN_TL_ axon extension to thoracic and lumbar segments, suggests that Cbln1 might function during CSN_TL_ axon extension. Further, the temporal restriction of *Cbln1* expression in layer V to early postnatal development suggests that the function of Cbln1 within CSN is potentially distinct from the function of Cbln1 in the cerebellum, where it is constitutively expressed throughout development and adulthood, and is required for both the proper formation and maintenance of synapses between Purkinje neurons and parallel fibers (Hirai et al., 2005).

### Other cerebellin family members are not expressed by CSN

Cbln1 forms homo- and hetero-complexes with itself or with other members of the Cbln family, and these complexes are known to perform varied, compensatory, and redundant functions (Pang et al., 2000a; Bao et al., 2006; Iijima et al., 2007; Miura et al., 2009; Joo et al., 2011; Rong et al., 2012; Seigneur et al., 2018; Seigneur and Südhof, 2018). To determine whether other *Cbln* family members are expressed by CSN_medial_ and might potentially interact with *Cbln1* in CSN_medial_, we examined previously published gene expression datasets and performed *in situ* hybridization in wild-type mice.

Differential gene expression analysis comparing CSN_BC-lat_ and CSN_medial_ indicates that *Cbln2, Cbln3*, and *Cbln4* are not expressed, or expressed at very low levels, by either CSN_BC-lat_ or CSN_medial_ during early postnatal development (Sahni et al., 2021a) (Fig. 4A,E,I). Accordingly, there is also no detectable difference between *Cbln2, Cbln3*, or *Cbln4* expression in CSN_BC-lat_ compared to CSN_medial_, suggesting that these genes are unlikely to function in a CSN subpopulation-specific manner. Interestingly, prior gene expression comparisons at late embryonic and early postnatal times found that *Cbln2* expression is enriched in a different neocortical projection neuron subtype, callosal projection neurons (CPN), as compared to CSN, and that *Cbln3* and *Cbln4* are not expressed by CPN or by CSN (Arlotta et al., 2005). Together, these datasets suggest that Cbln1 functions independently of other Cbln family members within CSN_medial_.

**Figure 4:**
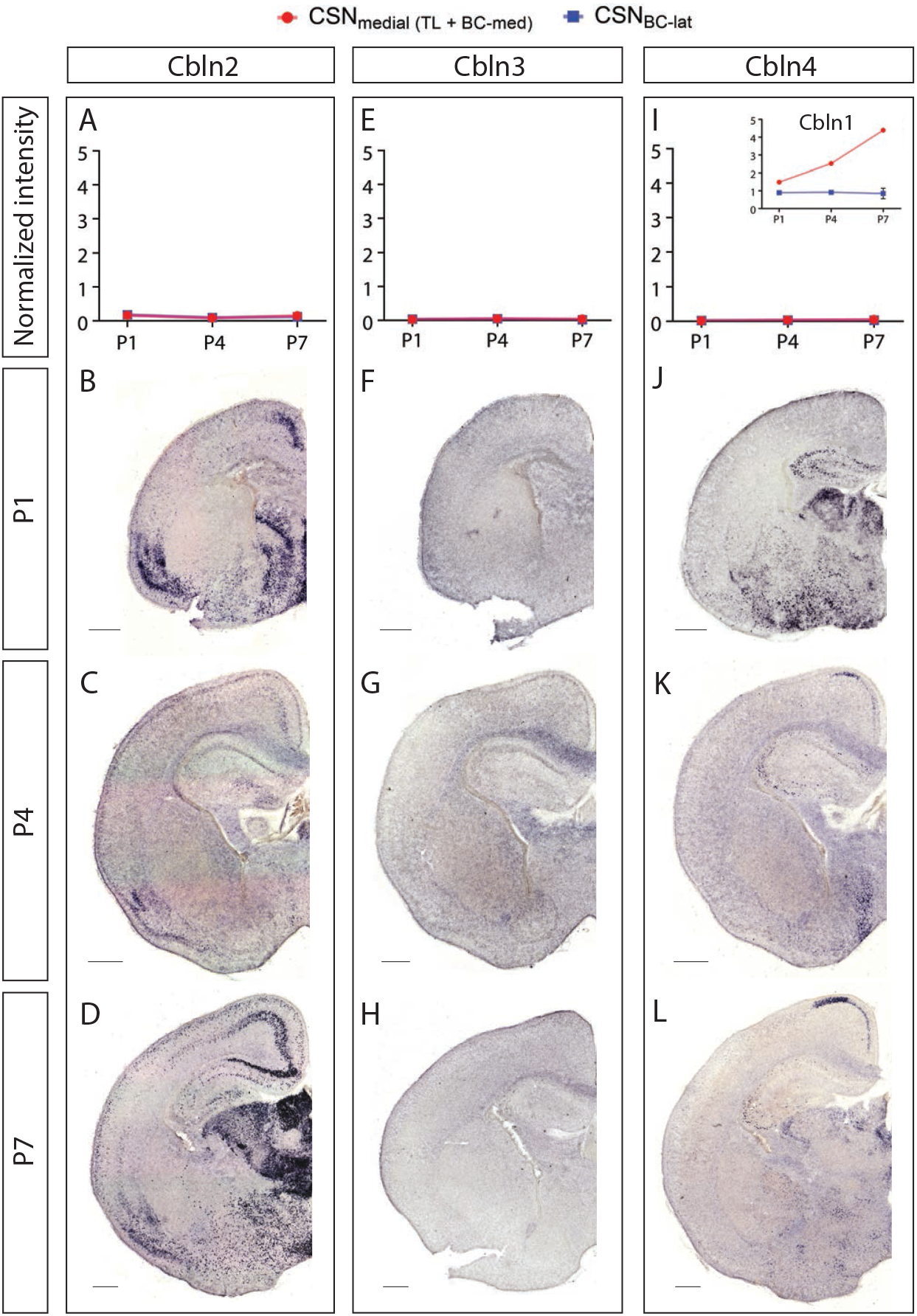
Other *Cbln* family members are not expressed by developing CSN. Prior differential gene expression analysis (Sahni et al., 2021a) identified *Cbln1* (inset in I) as expressed by CSN_medial_. In that dataset, *Cbln2* **(A)**, *Cbln3* **(E)**, and *Cbln4* **(I)** exhibit no expression by CSN_medial_ (blue) or CSN_BC-lat_ (red) at P1, P4, and P7. *In situ* hybridization at P1 (**B, F, J**), P4 (**C, G, K**), and P7 (**D, H, L**) confirms that all three genes are not expressed in layer V in the developing neocortex. Scale bars are 500μm.

*In situ* hybridization in wild-type mice confirmed the differential gene expression analyses. As reported previously (Miura et al., 2006; Seigneur and Südhof, 2017), *Cbln2* is expressed at high levels in cingulate and piriform cortex, and at intermediate levels in layer VI at P1, P4, and P7 (Fig. 4B-D). A small fraction of layer V neurons express *Cbln2* at P1 (Fig. 4B), but these are likely CPN based on the previous transcriptomic analysis (Arlotta et al., 2005). *Cbln3* is not expressed within sensorimotor cortex at P1, P4, or P7 (Fig. 4F-H). Similarly, *Cbln4* is not expressed in layer V, but is expressed in the cingulate at P4 and P7 (Fig. 4J-L). Together with the transcriptomic analyses, these results indicate that *Cbln2, Cbln3*, and *Cbln4* expression is likely absent in CSN, and thus, suggest that Cbln1 function in CSN_medial_ during early postnatal development is independent of other Cbln family members.

### Cbln1 is not required for CSN_TL_ axon extension to the thoracolumbar spinal cord

Since *Cbln1* expression levels coincide with the peak period of CSN_TL_ axon extension to thoracic and lumbar segments, we next investigated whether Cbln1 is necessary for CSN_TL_ axon extension to these distal spinal segments. We analyzed *Cbln1* null mice using anterograde and retrograde labeling (Hirai et al., 2005). We injected the retrograde tracer Cholera Toxin Subunit B conjugated with Alexa-Fluor 555 (CTB-555) into lumbar L1-L2 in *Cbln1* wild-type (WT), *Cbln1* heterozygous (Het), and *Cbln1* null mice at P5, at which time the first CSN_TL_ axons have reached the lumbar cord, and analyzed the number of retrogradely labeled CSN in sensorimotor cortex at P7 (Fig. 5A). As expected, labeled CSN are located caudomedially in sensorimotor cortex. We observe no overt difference in the overall distribution or number of retrogradely labeled CSN in *Cbln1* null mice (Fig. 5B-H).

**Figure 5:**
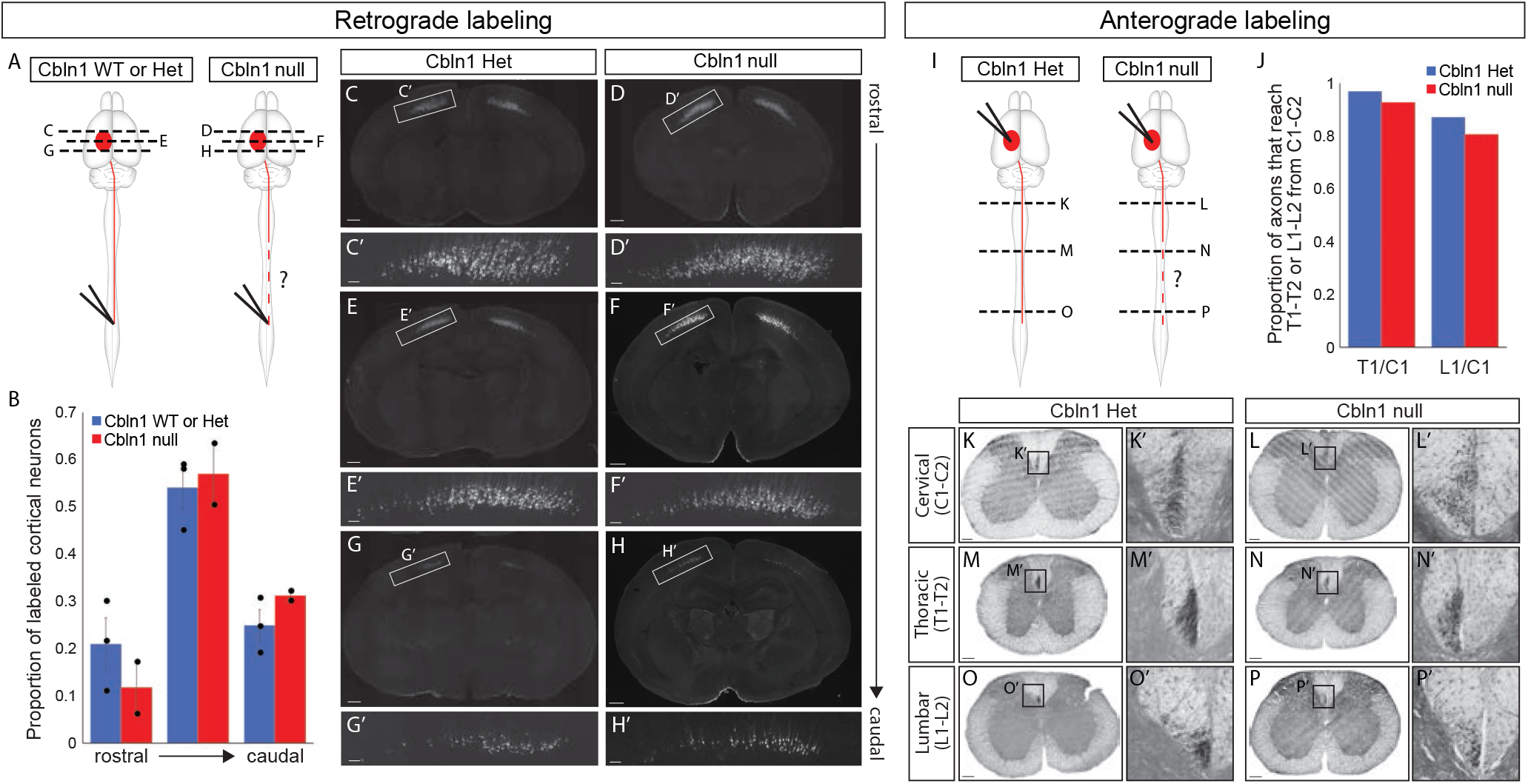
Cbln1 is not required for axon extension to lumbar L1-L2. **(A)** Experimental outline: Retrograde labeling with injection of CTB-555 at lumbar L1-L2 was performed in *Cbln1* WT, *Cbln1* Het, and *Cbln1* null mice at P5, and the number of retrogradely labeled neurons throughout the rostro-caudal extent of medial sensorimotor cortex was analyzed at P7. **(B)** There is no significant difference in the proportion of retrogradely labeled CSN in medial sensorimotor cortex at rostral (*p* = 0.34), middle (*p* = 0.73), or caudal (*p* = 0.24) levels by the two-tailed Student’s T-test between *Cbln1* WT or Het (*n* = 3) and *Cbln1* null (*n* = 2) mice. (**C**-**H**) Representative rostral, middle, and caudal sections from a *Cbln1* Het and a *Cbln1* null mouse. Scale bars are 100μm for insets and are 500μm for all other images. **(I)** Experimental outline: Anterograde labeling via biotinylated dextran amine (BDA) iontophoresis into caudomedial sensorimotor cortex was performed in *Cbln1* Het and *Cbln1* null mice at P28, and the number of BDA-labelled axons at cervical C1-C2, thoracic T1-T2, and lumbar L1-L2 was analyzed at P35. **(J)** There is no overt difference in the proportion of axons that reach thoracic T1-T2 or lumbar L1-L2 from cervical C1-C2 between *Cbln1* Het and *Cbln1* null mice. (**K**-**P**) Representative axial sections from cervical C1-C2, thoracic T1-T2, and lumbar L1-L2 in *Cbln1* Het and *Cbln1* null mice. Scale bars are 100μm.

Next, we injected the anterograde tracer biotinylated dextran amine (BDA) into the caudomedial sensorimotor cortex at P28 and analyzed the brain and spinal cord at P35 to determine whether *Cbln1* might be required for the maintenance of lumbar axon extension (Fig. 5I). Once again, there is no overt difference in the proportion of CSN axons at cervical C1-C2 that reach thoracic T1-T2 or lumbar L1-L2 between *Cbln1* WT and *Cbln1* null mice (Fig. 5J-P). These results suggest that *Cbln1* is not necessary for CSN_TL_ axon extension to the thoracic and lumbar cord.

### Mis-expression of Cbln1 in CSN_BC-lat_ leads to aberrant axon extension past the cervical cord toward distal thoraco-lumbar spinal segments

Although Cbln1 is not required for CSN_TL_ axon extension to distal thoraco-lumbar spinal segments, we wondered whether Cbln1 might be sufficient to direct long CSN axon extension. We had previously established that mis-expression of Crim1, a CSN_TL_-specific control, in CSN_BC-lat_ can redirect a subset of their axons to caudal thoracic segments (Sahni et al., 2021b). We therefore investigated whether Cbln1 is similarly sufficient to direct thoraco-lumbar axon extension by CSN_BC-lat_, which normally do not express *Cbln1*. To test this hypothesis, we introduced a plasmid expressing *Cbln1* and an EGFP reporter into CSN_BC-lat_ via *in utero* electroporation at E12.5. Control mice received a plasmid expressing EGFP alone (schematized in Fig. 6A). CSN_BC-lat_ axons normally reach the caudal cervical cord by P1 and never extend past the rostral-most segments in the thoracic cord (Sahni et al., 2021a). We examined electroporated mice at P4, by which time the most distally extending CSN_BC-lat_ axons have normally terminated within the caudal cervical or rostral-most segments of the thoracic cord.

**Figure 6:**
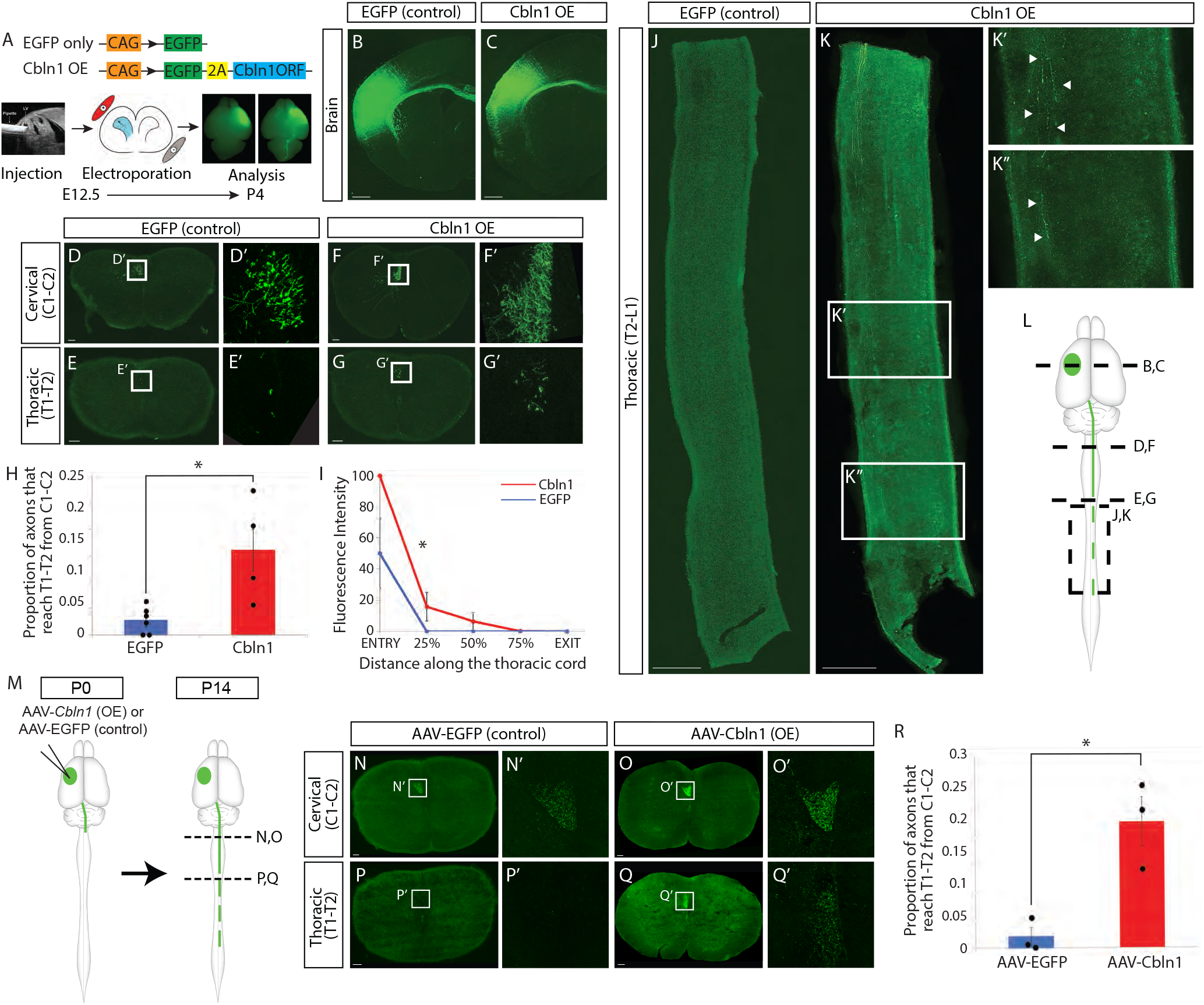
Cbln1 overexpression in CSN_BC-lat_ is sufficient to re-direct axon extension to distal thoracic spinal segments. **(A)** Experimental outline: In one set of experiments, plasmids were designed to express either *EGFP* alone, or both *EGFP* and *Cbln1* (Cbln1 OE). The constructs were delivered to developing CSN_BC-lat_ in lateral cortex using *in utero* electroporation at E12.5, and tissue was collected at P4 for analysis. CSN_BC-lat_ axons were visualized using immunocytochemistry for EGFP. (**B**,**C**) Electroporation location and distribution in the brain are well-matched between EGFP (control) and Cbln1 OE mice. (**D**-**G**) The number of CSN_BC-lat_ axons at cervical C1-C2 and thoracic T1-T2 were counted in axial sections of the spinal cord in EGFP and Cbln1 OE mice. **(H)** There are significantly more axons that reach thoracic T1-T2 from cervical C1-C2 in Cbln1 OE (red, *n* = 4) compared to EGFP (control; blue, *n* = 6) mice (*p* = 0.01 by two-tailed Student’s t-test). **(I)** CST fluorescence intensity was quantified along the rostro-caudal extent of the thoracic cord and normalized to the fluorescence intensity at thoracic T2. Significantly more *Cbln1*-expressing CSN_BC-lat_ axons extend into the distal thoracic cord when compared to EGFP controls (*p* = 0.04 by two-way ANOVA with repeated measures followed by Fisher’s least significant difference posthoc test). (**J**-**K**) CSN_BC-lat_ axons extend into the distal thoracic cord in thoracic sagittal sections in Cbln1 OE (arrowheads in K’, K”) but not in EGFP (control) mice. **(L)** Schematic indicating anatomical location of sections displayed in B-K. **(M)** Experimental outline: In a second set of experiments, AAV particles engineered to express either *EGFP* alone (AAV-EGFP) or both *EGFP* and *Cbln1* (AAV-Cbln1) were injected into rostrolateral sensorimotor cortex at P0 to test whether *Cbln1* is sufficient to re-direct axon extension by postmitotic CSN_BC-lat_. AAV-injected mice were then analyzed at P14. (**N**-**Q**) The number of axons that reach cervical C1-C2 and thoracic T1-T2 were counted in axial sections of the spinal cord in AAV-EGFP and AAV-Cbln1 mice. **(R)** There are significantly more axons that reach thoracic T1-T2 from cervical C1-C2 in AAV-Cbln1 (red, *n* = 3) compared to AAV-EGFP (blue, *n* = 3) (*p* = 0.02 by one-tailed Student’s t-test). Scale bars are 100μm for D-G and N-Q and 500μm for B, C, J, and K.

We first confirmed that all electroporations were restricted to lateral sensorimotor cortex where CSN_BC-lat_ reside (Fig. 6B,C). We next investigated the percentage of CSN_BC-lat_ axons that reach the rostral thoracic cord (T1-T2) from the rostral cervical cord (C1-C2). Strikingly, when Cbln1 is mis-expressed in CSN_BC-lat_, a 6-fold higher percentage of axons reach T1-T2 from C1-C2 (T1/C1) compared to the control (16.1% ± 4.0% for Cbln1 mis-expression, 2.8% ± 1.3% for the control; *p* = 0.01) (Fig. 6D-H). This indicates that Cbln1 mis-expression aberrantly directs a higher percentage of CSN_BC-lat_ axons to extend past their normal cervical targets into the thoracic cord by P4.

We next investigated whether these re-directed axons from *Cbln1*-expressing CSN_BC-lat_ extend further caudally toward more distal thoracic segments. As expected, in control mice, CSN_BC-lat_ axons do not extend past the rostral-most segments of the thoracic cord (Fig. 6J). In striking contrast, not only do a significantly larger number of CSN_BC-lat_ axons reach the rostral thoracic cord upon Cbln1 mis-expression, a subset of these axons reach the distal-most segments of the thoracic cord (T13) at P4 (Fig. 6K). Quantification of CST fluorescence intensity along the rostro-caudal extent of the thoracic cord (normalized to the fluorescence intensity at thoracic T2), reveals that significantly more Cbln1-expressing CSN_BC-lat_ axons extend into the distal thoracic cord when compared to controls (*p* = 0.04) (Fig. 6I). Indeed, no axons even reached T2 in 3 of the 6 control samples (Fig. 6I). Interestingly, although axon collateralization by CSN_BC-lat_ is well underway by P4 (Bareyre et al., 2005), we do not observe any collateralization by the aberrantly extended CSN_BC-lat_ axons upon Cbln1 mis-expression in the thoracic cord (Fig. 6K). Together, these data indicate that Cbln1 is sufficient to re-direct CSN_BC-lat_ axon extension past the cervical cord toward thoracic spinal segments but does not promote axon collateralization.

### Postnatal mis-expression of Cbln1 in CSN_BC-lat_ leads to aberrant long CSN_BC-lat_ axon extension

The time course of *Cbln1* expression suggests that its function is required specifically during the period of CSN_TL_ axon extension. However, since mis-expression by *in utero* electroporation begins in progenitors and continues into postmitotic neurons, there remained the unlikely possibility that the effect of Cbln1 mis-expression on CSN_BC-lat_ axon extension is due to alterations in early CSN_BC-lat_ specification before their axons reach the spinal cord, ultimately causing their aberrant axon extension later at P4. To directly investigate this possibility, we performed mis-expression in CSN_BC-lat_ at P0 via AAV-mediated gene delivery. We generated AAV particles engineered to express Cbln1 and an EGFP reporter (AAV-Cbln1). Control mice received AAV particles engineered to express EGFP alone (AAV-EGFP). We injected these particles into the rostrolateral cortex of P0 mice under ultrasound guided backscatter microscopy and examined CSN_BC-lat_ axon extension at P14 (schematized in Fig. 6M), at which point CSN axonal projections have been pruned and the adult connectivity pattern of the CST is largely established (Bareyre et al., 2005; Sahni et al., 2020, 2021a).

As with embryonic mis-expression of Cbln1 in CSN_BC-lat_, postnatal Cbln1 mis-expression leads to aberrant CSN_BC-lat_ axon extension past the cervical cord into the thoracic spinal cord. We quantified the number of axons that reach thoracic T1-T2 compared to the number of axons at cervical C1-C2 in axial sections from mice injected with either AAV-EGFP or AAV-Cbln1 (Fig. 6N-R). There is a significant increase in the percentage of axons that reach thoracic T1-T2 at P14 compared to controls (22.8% ± 4.5% for Cbln1 mis-expression, 2.0% ± 1.7% for the control; *p* = 0.02) (Fig. 6R). As with the *in utero* electroporation experiments, these aberrantly extended CSN_BC-lat_ axons upon Cbln1 mis-expression at P0 also fail to collateralize in the thoracic cord (data not shown). Therefore, postnatal mis-expression of Cbln1 at P0 is sufficient to direct CSN_BC-lat_ axons past the cervical cord and into the thoracic cord. Together, these data combined with the results from *in utero* mis-expression experiments indicate that Cbln1 regulates axon extension without affecting CSN axon collateralization. This is in contrast to known functions of Cbln1 as a critical synaptic organizer in the cerebellum, striatum, and other brain regions (Hirai et al., 2005; Kusnoor et al., 2010; Seigneur and Südhof, 2018), and represents a unique function of Cbln1 in controlling CSN axon extension independent of effects on axon collateralization or synapse formation.

### Postnatal overexpression of Cbln1 in CSN_medial_ is sufficient to drive long CSN axon extension to the thoraco-lumbar spinal cord

The experiments above indicate that CSN_TL_ axon extension to caudal thoracic and lumbar segments does not require Cbln1 function, but that Cbln1 is sufficient to re-direct CSN_BC-lat_ axons to the caudal thoracic cord. Unlike CSN in lateral sensorimotor cortex, which only project to bulbar-cervical segments (CSN_BC-lat_), CSN_medial_ include distinct subpopulations with projections to bulbar-cervical (CSN_BC-med_) or thoracolumbar segments (CSN_TL_). Previous work has established that CSN_BC-lat_, CSN_BC-med_, and CSN_TL_ are distinct, molecularly delineated subpopulations that can be distinguished before CSN axons have even reached the spinal cord (schematized in Fig. 1A) (Sahni et al., 2021a).

To determine whether Cbln1 controls axon extension past the cervical cord by multiple molecularly and spatially distinct subpopulations, we next investigated whether Cbln1 overexpression is sufficient to re-direct CSN_BC-med_ axons toward caudal thoracic and lumbar segments. We overexpressed Cbln1 at P0 via AAV-mediated gene delivery in medial sensorimotor cortex, where *Cbln1* is normally expressed by only a subset of CSN_medial_. We injected the medial cortex of P0 mice with either AAV particles engineered to express Cbln1 and an EGFP reporter (AAV-Cbln1) or AAV particles engineered to express EGFP alone (AAV-EGFP). We examined CSN_medial_ axon extension at P14 by quantifying CST fluorescence intensity at thoracic T1-T2 and lumbar L1-L2, normalized to the fluorescence intensity at cervical C1-C2 in axial sections from mice injected with either AAV-Cbln1 or AAV-EGFP (schematized in Fig. 7A).

**Figure 7:**
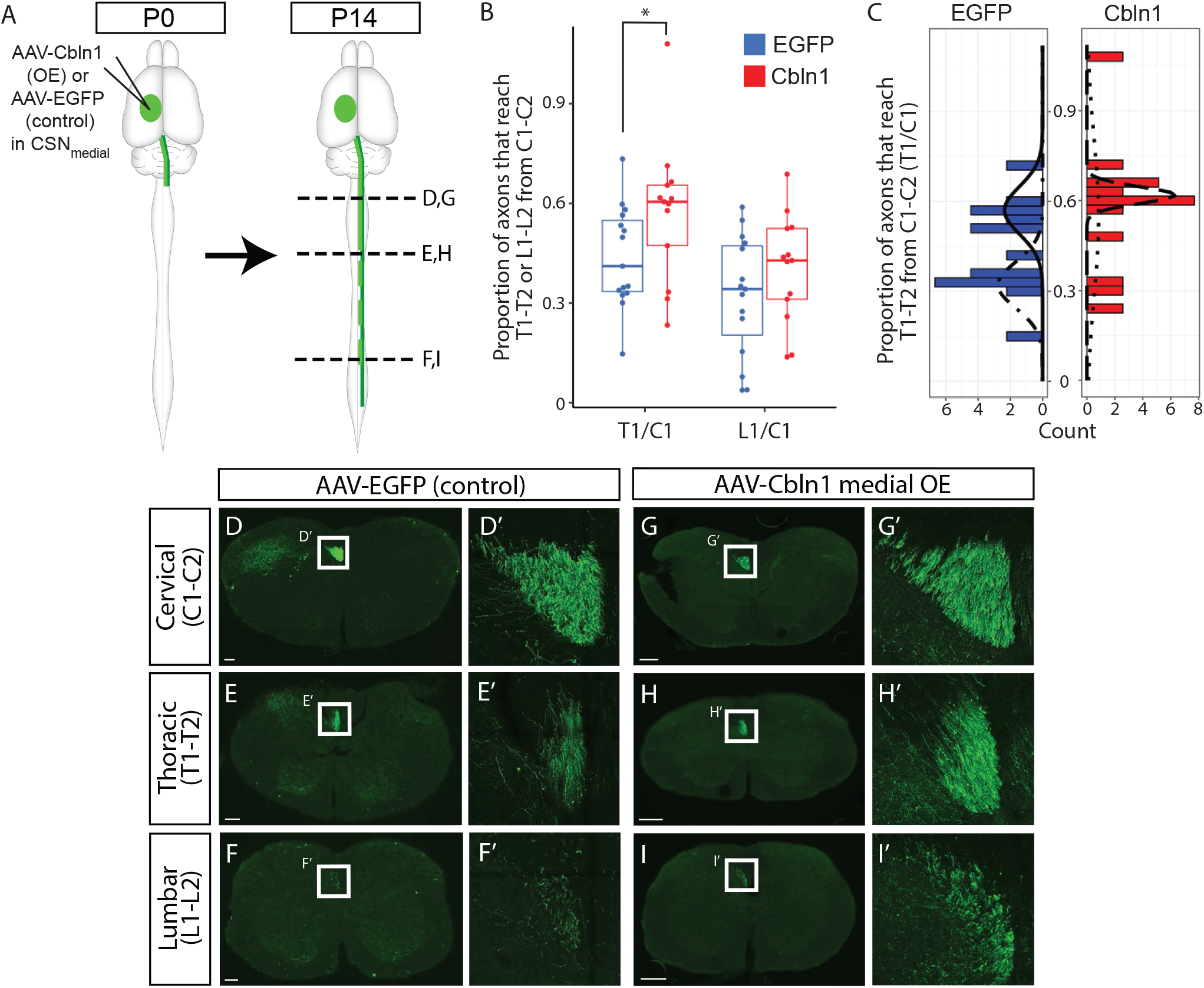
Cbln1 overexpression in CSN_medial_ is sufficient to increase the number of axons extending past the cervical spinal cord. **(A)** Experimental outline: AAV particles engineered to express either *EGFP* alone (AAV-EGFP, control) or both *EGFP* and *Cbln1* (AAV-Cbln1) were injected into medial sensorimotor cortex at P0. AAV-injected mice were analyzed at P14. **(B)** CST intensity was quantified in axial sections at cervical C1-C2, thoracic T1-T2, and lumbar L1-L2. The proportion of axons that reach thoracic T1-T2 from cervical C1-C2 (T1/C1) is significantly higher in AAV-Cbln1 (red, *n* = 13) compared to AAV-EGFP (blue, *n* = 15) (*p* = 0.03 by one-tailed Student’s t-test). In contrast, the proportion of axons that reach lumbar L1-L2 from cervical C1-C2 is not significantly different between AAV-Cbln1 and AAV-EGFP mice (*p* = 0.11 by one-tailed Student’s t-test). **(C)** We modeled the distribution of T1/C1 in control AAV-EGFP or AAV-Cbln1 injected mice as a mixture of two Gaussians. The distribution of T1/C1 in control AAV-EGFP injected mice appears bimodal with one Gaussian centered at 0.32 ± 0.08 and the other at 0.57 ± 0.08, likely reflecting variability in the proportion of CSN_BC-med_ or CSN_TL_ labeled by each injection. In contrast, the distribution of T1/C1 in AAV-Cbln1 injected mice appears unimodal with one Gaussian centered at 0.54 ± 0.27 and the other centered at 0.62 ± 0.03. These Gaussians are similar to the Gaussian with control AAV-EGFP injections centered at 0.57, which is likely comprised of a higher proportion of CSN_TL_ compared to CSN_BC-med_. This suggests that Cbln1 overexpression might specifically shift the segmental targeting of CSN_BC-med_ past the bulbar-cervical cord into the thoraco-lumbar cord, but not overtly affect the segmental targeting of CSN_TL_. (**D**-**I**) Representative axial sections from cervical C1-C2, thoracic T1-T2, and lumbar L1-L2 from control AAV-EGFP and AAV-Cbln1 injected mice. Scale bars are 100μm.

Since there is no positive molecular identifier for CSN_BC-med_, we investigated their axon targeting as a subset of the broader CSN_medial_ subpopulation. Given the two-population diversity of CSN_medial_, injections in medial sensorimotor cortex will label varying proportions of CSN_BC-med_ and CSN_TL_, since both subpopulations reside interdigitated in medial sensorimotor cortex. Therefore, we expected these injections to result in greater variability in the proportion of labeled CSN_medial_ axons that reach thoracic T1-T2 from cervical C1-C2 (we refer to this ratio as T1/C1). Indeed, we observe high variance in the distribution of T1/C1 following injection with control AAV-EGFP (Fig. 7B). Importantly, the AAV-EGFP-injected mice clustered broadly into two groups, likely reflecting the predicted differences in the extent of labeling between CSN_BC-med_ and CSN_TL_ in each individual mouse. Therefore, we modeled the distribution of T1/C1 as a mixture of two Gaussians (Fig. 7C). The distribution of T1/C1 following the injection of control AAV-EGFP appears bimodal, with one Gaussian centered at 0.32 ± 0.08– likely reflecting a higher proportion of CSN_BC-med_ relative to CSN_TL_ in these injections– and the other Gaussian at 0.57 ± 0.08– likely reflecting a higher proportion of CSN_TL_ relative to CSN_BC-med_.

Despite these differences in the relative numbers of CSN_BC-med_ and CSN_TL_ labeled by each injection, we investigated whether Cbln1 overexpression is sufficient to alter the T1/C1 ratio in AAV-Cbln1-injected mice. As with misexpression of Cbln1 in CSN_BC-lat_, we find that overexpression of Cbln1 in CSN_medial_ leads to long CSN axon extension past the cervical cord to thoraco-lumbar spinal segments (Fig. 7B-I). There is a significant increase in T1/C1 (57.5% ± 5.9% for Cbln1 overexpression, 43.8% ± 3.9% for the control; *p* = 0.03). Intriguingly, we do not find a similar increase in the proportion of axons that reach lumbar L1-L2 from cervical C1-C2 (40.3% ± 4.5% for Cbln1 overexpression, 32.1% ± 4.7% for the control; *p* = 0.11) upon Cbln1 overexpression in CSN_medial_. In contrast with the T1/C1 distribution in control AAV-EGFP-injected mice, which appears bimodal, the distribution of T1/C1 following Cbln1 overexpression appears unimodal with the center of both Gaussian distributions around 0.57 (Fig. 7C). This is similar to the Gaussian distribution in the control T1/C1 distribution that is likely enriched for CSN_TL_. Although this analysis does not entail *a priori* cell identification, it suggests that Cbln1 overexpression might specifically shift the segmental targeting of CSN_BC-med_ past the bulbar-cervical cord into the thoraco-lumbar cord, but might not substantially affect the segmental targeting of CSN_TL_. Together with our findings that Cbln1 mis-expression in CSN_BC-lat_ promotes axon extension past their normal cervical targets, these results indicate that Cbln1 is sufficient to drive axon extension past the cervical cord by multiple spatially and molecularly distinct CSN subpopulations.

### Cbln1 is regulated by Klhl14, but acts independently of Crim1 to control CSN long axon extension

We previously identified Klhl14 and Crim1 as molecular controls expressed by CSN_BC-lat_ and CSN_TL_, respectively (Sahni et al., 2021a,b). Klhl14 functions to limit CSN_BC-lat_ axons to proximal segments in the cervical spinal cord during early postnatal development. Knockdown of *Klhl14* by shRNA leads to aberrant CSN_BC-lat_ axon extension toward distal thoracic segments at P4. This aberrant axon extension is accompanied by an upregulation of *Crim1* by CSN_BC-lat_.

To determine whether Klhl14 might also similarly modulate *Cbln1* expression, we examined coronal brain sections from mice at P4 in which *Klhl14* shRNA was introduced into CSN_BC-lat_ via *in utero* electroporation at E12.5 (Fig. 8A). Strikingly, *Cbln1* is ectopically expressed in layer V in rostrolateral sensorimotor cortex where *Klhl14* is reduced, but not in the contralateral cortex where *Klhl14* expression is normal (Fig. 8B). This strongly suggests that *Cbln1* and *Crim1* expression are both repressed by Klhl14 in CSN_BC-lat_ to regulate CSN axon extension.

**Figure 8:**
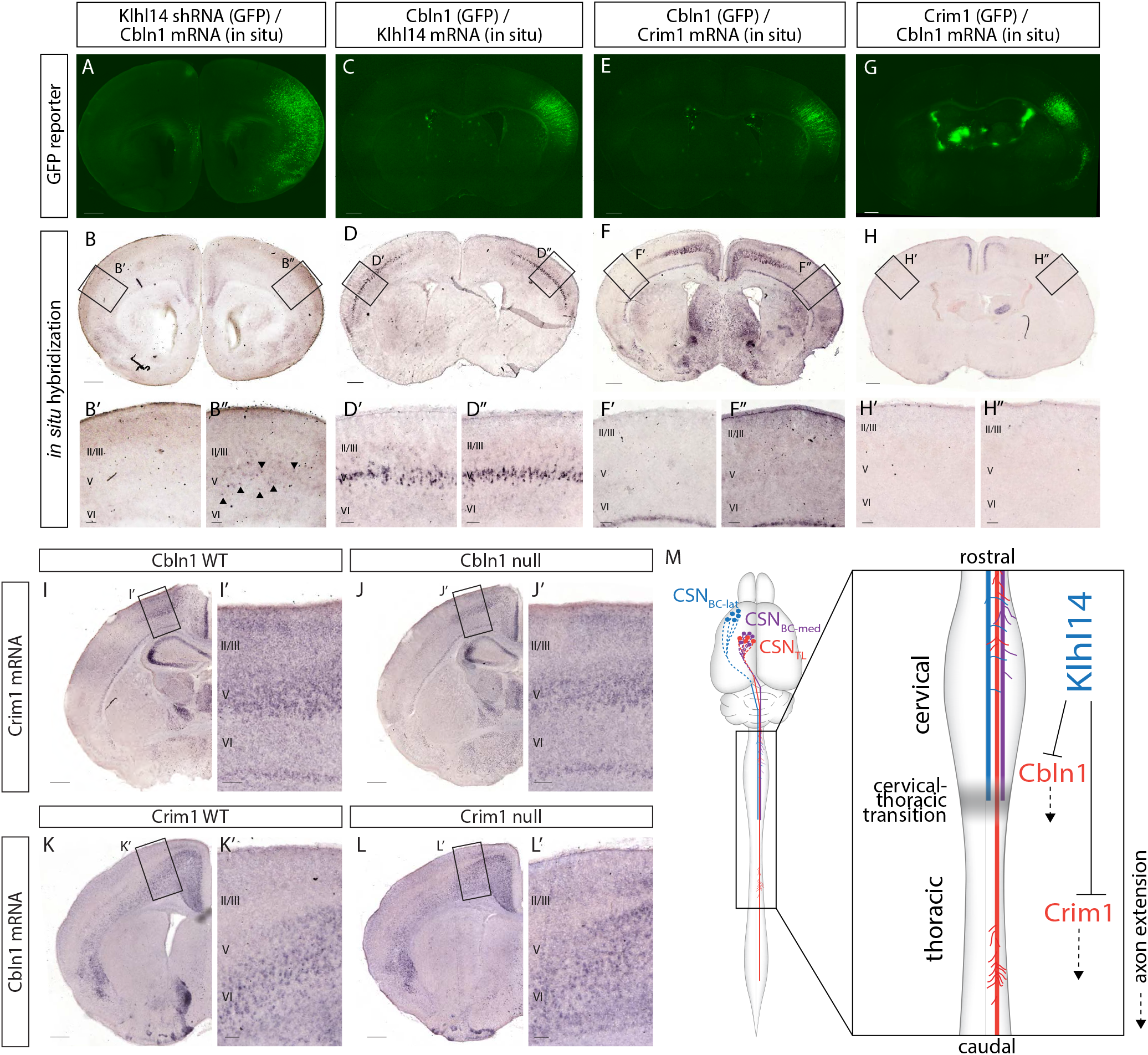
*Cbln1* expression is regulated by Klhl14 but not by Crim1. (**A**,**B**) Coronal section of a P4 brain that was electroporated *in utero* at E12.5 with Klhl14 shRNA. **(A)** EGFP fluorescence (green) shows the site of electroporation in lateral cortex. **(B)** *In situ* hybridization image of the same section in (A) shows *Cbln1* expression. *Cbln1* is normally restricted to medial cortex. However, *Klhl14* knockdown by shRNA causes ectopic *Cbln1* expression in lateral cortex (arrowheads in B”) in the electroporated cortical hemisphere (compare B” with contralateral B’). (**C**,**D**) Coronal section of a P4 brain that was electroporated *in utero* at E12.5 with a plasmid containing *Cbln1* and *EGFP* (Cbln1-EGFP). **(C)** EGFP fluorescence (green) shows the site of electroporation in lateral cortex. **(D)** *In situ* hybridization image of the same section in (C) showing *Klhl14* expression in Cbln1-misexpressing CSN_BC-lat_ remains unchanged, indicating that *Klhl14* expression in lateral layer V is unaffected by Cbln1 mis-expression. (**E**,**F**) Coronal section of a P4 brain that was electroporated *in utero* at E12.5 with Cbln1-EGFP. **(E)** EGFP fluorescence (green) shows the site of electroporation in lateral cortex. **(F)** *In situ* hybridization image of the same section in (E) showing that there is no ectopic *Crim1* expression in Cbln1-misexpressing CSN_BC-lat_. (**G**,**H**) Coronal section of a P4 brain that was electroporated *in utero* at E12.5 with Crim1-EGFP. *Crim1* mis-expression in CSN_BC-lat_ can redirect axons toward caudal thoracic spinal segments (Sahni et al., 2021b). **(G)** EGFP fluorescence (green) shows the site of electroporation in lateral cortex. **(H)** *In situ* hybridization image of the same section in (G) showing that there is no ectopic *Cbln1* expression in Crim1misexpressing CSN_BC-lat_. (**I**,**J**) *Crim1* expression in medial layer V does not differ between *Cbln1* WT and *Cbln1* null mice. (**K**,**L**) *Cbln1* expression in medial layer V does not differ between *Crim1* WT and *Crim1* null mice. **(M)** Summary schematic displaying molecular controls over CSN axon extension both at, and beyond, the transition between cervical and thoracic spinal segments. Together with the previous investigations identifying Crim1 and Klhl14 function (Sahni et al., 2021b), our data suggest a model whereby Cbln1 directs CSN axon extension from the cervical into the thoracic cord, whereas Crim1 directs those CSN axons that cross this transition zone to extend further toward caudal thoracic and lumbar spinal segments. This indicates that CSN segmental axon targeting toward thoracic and lumbar segments involves multiple, distinct molecular regulators acting at distinct spinal levels. *Klhl14*, which is specifically expressed in CSN_BC-lat_ and restricts CSN_BC-lat_ axon extension to the bulbar-cervical segments, represses the expression of both *Cbln1* and *Crim1* in CSN_BC-lat_. This indicates that Klhl14 represses a broad program of thoraco-lumbar directed axon extension in CSN_BC-lat_. This program, mediated by multiple independent molecular controls, would otherwise direct CSN axons past the cervical cord toward caudal thoracic and lumbar segments. Scale bars are 100μm for insets and 500μm for all other images.

We also examined whether Cbln1 overexpression can regulate *Klhl14* expression. Coronal brain sections from mice electroporated with *Cbln1* in lateral cortex at E12.5 and analyzed at P4 were examined for *Klhl14* expression via *in situ* hybridization (Fig. 8C). There is no difference in *Klhl14* expression in the electroporated versus contralateral cortex (Fig. 8D). This strongly suggests that Klhl14 acts upstream of *Cbln1* to repress *Cbln1* expression by CSN_BC-lat_.

Gain-of-function experiments show that Crim1 and Cbln1 function at distinct levels of the spinal cord. Crim1 misexpression does not increase the proportion of CSN_BC-lat_ axons that reach thoracic T1-T2, but it does re-direct the small minority of CSN_BC-lat_ axons that reach the thoracic cord to extend farther into the thoracic cord toward caudal thoracic segments (Sahni et al., 2021b). In contrast, Cbln1 mis-expression significantly increases axon extension to thoracic T1-T2 by both CSN_BC-lat_ and CSN_medial_. These data suggest that Cbln1 and Crim1 function in distinct pathways, and that Klhl14 acts as an upstream regulator of both *Cbln1* and *Crim1*.

These results led us to investigate whether *Cbln1* and *Crim1* control CSN long axon extension via the same or distinct genetic pathways. We first examined *Crim1* expression in sensorimotor cortex in *Cbln1* WT and *Cbln1* null mice. We detect no differences in *Crim1* expression (Fig. 8I,J). Similarly, we detect no difference in *Cbln1* expression in sensorimotor cortex between *Crim1* WT and *Crim1* null mice (Fig. 8K,L). We further investigated whether Cbln1 mis-expression in CSN_BC-lat_ via *in utero* electroporation at E12.5 might modulate *Crim1* expression when analyzed at P4, and vice versa. There is no difference in *Crim1* expression between *Cbln1*-expressing CSN_BC-lat_ compared to the contralateral cortex (Fig. 8E,F). There is also no difference in *Cbln1* expression between *Crim1*-expressing CSN_BC-lat_ and the contralateral cortex (Fig. 8G,H). Together, these data indicate that *Crim1* and *Cbln1* act via distinct genetic pathways to control CSN axon extension.

## Discussion

Previous work has identified that CSN subpopulations exhibit striking axon extension specificity during development, and that this specificity is durably maintained into maturity (Sahni et al., 2021a,b). CSN subpopulations with distinct spinal segmental targets are molecularly distinct from the earliest stages of axon extension, even before their axons reach the spinal cord. We previously investigated two molecular controls– Klhl14 and Crim1– that both prospectively identify CSN subpopulations with segmentally distinct projections, and control these projections. We identified their critical functions in directing CSN axons to appropriate spinal segmental levels, with dual-directional, complementary regulation toward thoraco-lumbar extension (by Crim1) and limiting axon extension past bulbar-cervical segments (by Klhl14). These results indicate that CSN-intrinsic molecular controls, at least in part, govern CSN axonal targeting specificity (Sahni et al., 2021a,b). Here, we build on this work to identify a novel role for a member of the cerebellin family, Cbln1, in controlling CSN segmental axonal projection targeting. We find that Cbln1 is expressed specifically by CSN in medial sensorimotor cortex. The time course of Cbln1 expression by CSN closely aligns with the period of CSN_TL_ axon extension to thoracic and lumbar segments. Mis-expression of Cbln1 in CSN_BC-lat_ via either *in utero* electroporation at E12.5 or AAV injection at P0 re-directs CSN_BC-lat_ axons past their normal cervical targets to distal segments in the thoraco-lumbar cord. Similarly, Cbln1 overexpression in CSN_medial_ is sufficient to increase the proportion of CSN_medial_ axons that extend to thoracic spinal segments. These results indicate that Cbln1 can direct long axon extension by multiple CSN subpopulations residing in spatially distinct locations in sensorimotor cortex.

This represents a novel function for Cbln1 in axon extension, independent of its well-described function as a synaptic organizer. Cbln1 has been characterized extensively in the cerebellum, where it is localized and secreted at presynaptic terminals of cerebellar granule cells, and is instructive for synapse formation between Purkinje cells and the parallel fibers of granule cells (Hirai et al., 2005; Matsuda et al., 2010; Uemura et al., 2010; Ibata et al., 2019). Outside of the cerebellum, Cbln1 has also been shown to play critical roles in both synapse formation and synapse maintenance in a number of other brain regions, including the hippocampus, striatum, and the ventral tegmental area of the midbrain (Kusnoor et al., 2010; Krishnan et al., 2017; Seigneur and Südhof, 2018).

Strikingly, we find that its function in CSN is quite distinct. *Cbln1* expression by CSN is strongest during the time period of axon extension, prior to axon collateralization or synapse formation. Moreover, although Cbln1 mis-expression in CSN_BC-lat_ re-directs these axons to the thoracic cord, these redirected axons do not collateralize in the thoracic cord at either P4 or P14. This suggests that the function and mechanism of Cbln1 in axon extension occurs independent of synapse formation by CSN. This function is likely directly mediated by Cbln1 localized to CSN axons; Cbln1 has been shown repeatedly to be localized at presynaptic terminals to directly affect synapse formation and maintenance in multiple brain regions (Matsuda et al., 2010; Otsuka et al., 2016), and is likely trafficked similarly along CSN axons to directly regulate axon extension. Our data intriguingly suggest that Cbln1 and other classical synaptic organizers might perform currently unappreciated functions in axon extension during development.

Further work will be required to identify proteins with which Cbln1 interacts to mediate its function in CSN axon extension. Cbln1 interactors such as GluRδ2 and β-neurexins have been identified in the cerebellum (Matsuda et al., 2010; Uemura et al., 2010; Elegheert et al., 2016; Cheng et al., 2016), and there is evidence that, in the cerebral cortex, where GluRδ2 is not expressed, Cbln1 instead interacts with GluRδ1 (Matsuda et al., 2010; Rong et al., 2012; Wei et al., 2012; Yasumura et al., 2012). *GRID1* and *GRID2* (which encode GluRδ1 and GluRδ2, respectively), as well as neurexin family members, are expressed in the human spinal cord (GTEx Consortium, 2013). These interactors have not been implicated previously in axon extension; however, it is possible that they might also interact with Cbln1 to promote long axon extension in this novel context.

Although Cbln1 overexpression robustly directs long CSN axon extension, we do not observe any substantial defects of CSN_TL_ axon extension to thoracic and lumbar spinal segments in *Cbln1* null mice, suggesting that Cbln1 is sufficient but not necessary for CSN long axon extension. What might compensate for Cbln1 function in CSN_TL_ in *Cbln1* null mice? The other cerebellin family members, *Cbln2, Cbln3*, and *Cbln4*, are known to interact with *Cbln1* and perform compensatory or redundant functions (Bao et al., 2006; Iijima et al., 2007; Miura et al., 2009; Pang et al., 2000a; Joo et al., 2011; Rong et al., 2012; Seigneur et al., 2018; Seigneur and Südhof, 2018). For instance, *Cbln1* and *Cbln2* are normally expressed in the cerebellum. Whereas *Cbln1* null mice are ataxic and have disrupted synaptic connectivity, *Cbln2* null mice display no functional or anatomical deficits in the cerebellum. Interestingly, however, Cbln2 over-expression in the cerebellum of *Cbln1* null mice can rescue their ataxic phenotype, suggesting partial redundancy of function (Rong et al., 2012). Although compensation by other cerebellins presents a tempting hypothesis, we find that *Cbln2, Cbln3*, and *Cbln4* are not normally expressed by CSN_medial_ at the time when their axons are normally extending to distal spinal segments. It is possible that molecular controls other than Cbln protein family members might compensate for loss of Cbln1 in CSN_medial_. For instance, Cbln1 and neuroligin3 have been shown to partially compensate for each other at calyx of Held synapses (Zhang et al., 2017; Yuzaki, 2017). Future studies will be needed to elucidate molecular mechanisms that might compensate for the loss of Cbln1 function in directing CSN axon extension.

We also investigated whether Cbln1 might regulate or be regulated by previously identified molecular controls over CSN axon extension. We previously identified that Klhl14, which is specifically expressed by CSN_BC-lat_ and limits their axon extension to the cervical cord, acts at least in part to repress *Crim1* expression by CSN_BC-lat_. Crim1 mis-expression in CSN_BC-lat_ is sufficient to re-direct their axons to distal thoracic levels (Sahni et al., 2021b). In the work presented here, we find that Klhl14 also represses *Cbln1* expression by CSN_BC-lat_, suggesting that Klhl14 acts as a broad transcriptional repressor to suppress multiple molecular controls that otherwise would direct CSN axons past the cervical cord. However, we find that *Cbln1* and *Crim1* are not in the same genetic pathway in CSN. In either *Cbln1* null or *Crim1* null mice, there is no change in the expression of the other molecular control. Likewise, when either Cbln1 or Crim1 is misexpressed in CSN_BC-lat_, the other molecular control is not ectopically expressed.

Indeed, Cbln1 and Crim1 perform similar, but distinct, functions in directing long CSN axon extension. Mis-expression of Cbln1, but not Crim1 (Sahni et al., 2021b), in CSN_BC-lat_ increases the number of axons that reach thoracic T1-T2. Further, although both Cbln1 and Crim1 mis-expression in CSN_BC-lat_ is sufficient to re-direct those axons that reach T1-T2 to extend further into the thoracic cord, their effects on axon extension within the thoracic cord are distinct. While 100% of mice in which Crim1 was misexpressed in CSN_BC-lat_ had axons that reached at least halfway through the thoracic cord (Sahni et al., 2021b), this was true of only 50% of mice in which Cbln1 was misexpressed in lateral cortex. This suggests that Cbln1 might serve as a regulator of axon targeting at the transition between cervical and thoracic spinal segments, whereas *Crim1* primarily functions to drive axon extension distal to this transition (schematized in Fig. 8M).

Additionally, CSN_BC-lat_ axons that aberrantly extend in the thoracic cord upon either *Cbln1* or *Crim1* mis-expression fail to collateralize in the thoracic spinal gray matter. It is likely that distinct molecular controls are required to interact with extracellular cues specific to the thoraco-lumbar spinal cord to promote axon collateralization. This is consistent with previous indications that axon extension and collateralization are distinctly regulated processes (Kalil and Dent, 2014; Itoh et al., 2021).

Interestingly and potentially relevant, motor neuron diseases (MND) such as amyotrophic lateral sclerosis (ALS) and hereditary spastic paraplegia (HSP) do not affect all CSN equally (Bruijn et al., 2004; Strong and Gordon, 2005; Salinas et al., 2008). In bulbar forms of ALS, e.g., brainstem-projecting CSN are affected, while in HSP, lumbar-projecting CSN preferentially degenerate. Although multiple proteins including SOD1 and TDP-43 have been implicated in MND (Hardiman et al., 2017), it remains unclear why certain CSN subpopulations preferentially degenerate in distinct MND subtypes. Identification and characterization of molecular controls that govern axonal development and connectivity of specific CSN subpopulations might provide insight regarding molecular mechanisms underlying preferential vulnerability of specific CSN subpopulations to degeneration.

Finally, our results suggest that Cbln1 might be a relevant molecular control for potential application in spinal cord repair and/or regeneration, and might elucidate broader organizing principles for establishing diverse connectivity by other neocortical projection neuron subtypes. Reactivating developmental controls to regulate CSN axon extension and sprouting might offer a promising approach to re-establish with some specificity damaged connectivity following spinal cord injury. More broadly, identification of molecular controls over development of anatomically and functionally diverse CSN subpopulations might elucidate fundamental principles of evolutionary diversification within originally more homogeneous neuronal populations and circuitry, while also offering potentially novel avenues for regeneration and/or repair of diseased and/or damaged neocortical or other nervous system circuitry.

## Acknowledgements

We thank Julia Kaiser for designing the schematics first shown in Figure 1A. We thank members of the Macklis and Sahni Labs for useful discussions and comments on the manuscript. This research was supported by National Institutes of Health (NIH) R01 NS045523, the Emily and Robert Pearlstein Fund, and a grant from the Travis Roy Foundation to J.D.M., NIH (NTRAIN/NICHD K12HD093427), and grants from the Wings for Life - Spinal Cord Injury Foundation and the Craig Neilsen Foundation, as well as additional infrastructure support from the Burke Foundation to V.S.. C.R. was partially supported by a postdoctoral fellowship from the Spinal Cord Injury Trust Fund through New York State Department of Health (SCIRB C33613GG). Additional infrastructure support from NIH DP1 NS106665, NS075672, Max and Anne Wien Professor of Life Sciences fund, and support from the DEARS Foundation to J.D.M.

